# Rapid blood acid-base regulation by European sea bass (*Dicentrarchus labrax*) in response to sudden exposure to high environmental CO_2_

**DOI:** 10.1101/2021.04.22.441026

**Authors:** Daniel W. Montgomery, Garfield T. Kwan, William G. Davison, Jennifer Finlay, Alex Berry, Stephen D. Simpson, Georg H. Engelhard, Silvana N.R. Birchenough, Martin Tresguerres, Rod W. Wilson

## Abstract

Fish in coastal ecosystems can be exposed to acute variations in CO_2_ that can approach 1 kPa CO_2_ (10,000 μatm). Coping with this environmental challenge will depend on the ability to rapidly compensate the internal acid-base disturbance caused by sudden exposure to high environmental CO_2_ (blood and tissue acidosis); however, studies about the speed of acid-base regulatory responses in marine fish are scarce. We observed that upon exposure to ~1 kPa CO_2_, European sea bass (*Dicentrarchus labrax*) completely regulate erythrocyte intracellular pH within ~40 minutes, thus restoring haemoglobin-O_2_ affinity to pre-exposure levels. Moreover, blood pH returned to normal levels within ~2 hours, which is one of the fastest acid-base recoveries documented in any fish. This was achieved via a large upregulation of net acid excretion and accumulation of HCO_3_− in blood, which increased from ~4 to ~22 mM. While the abundance and intracellular localisation of gill Na^+^/K^+^-ATPase (NKA) and Na^+^/H^+^ exchanger 3 (NHE3) remained unchanged, the apical surface area of acid-excreting gill ionocytes doubled. This constitutes a novel mechanism for rapidly increasing acid excretion during sudden blood acidosis. Rapid acid-base regulation was completely prevented when the same high CO_2_ exposure occurred in seawater with experimentally reduced HCO_3_− and pH, likely because reduced environmental pH inhibited gill H^+^ excretion via NHE3. The rapid and robust acid-base regulatory responses identified will enable European sea bass to maintain physiological performance during large and sudden CO_2_ fluctuations that naturally occur in coastal environments.

**Summary statement:** European sea bass exposed to 1 kPa (10,000 μatm) CO_2_ regulate blood and red cell pH within 2 hours and 40 minutes, respectively, protecting O_2_ transport capacity, via enhanced gill acid excretion.

## Introduction

Increased CO_2_ in aquatic environments, or environmental hypercapnia, causes significant physiological challenges for water breathing animals including fish. As environmental CO_2_ increases, there is a corresponding rise in CO_2_ within the fish’s blood, which in turn induces a decrease in blood pH. This condition is referred to as a respiratory acidosis, and depending on its magnitude, can disrupt multiple homeostatic processes including gas exchange (Crocker and Cech Jr, 1998; Eddy et al., 1977; Perry and Kinkead, 1989) and cardiovascular function (Lee et al., 2003; Perry and McKendry, 2001; Perry et al., 1999). Globally, CO_2_ levels in the ocean are increasing as a result of anthropogenic climate change, and are predicted to reach ~0.1 kPa (0.1% CO_2_, 1,000 μatm) by 2100 under a ‘business as usual’ scenario (Orr et al., 2005; Doney et al., 2011; IPCC, 2014).

The increase in oceanic CO_2_ levels, known as ocean acidification (OA) (Doney et al., 2009), has renewed interest in acid-base regulatory mechanisms of aquatic organisms. However, coastal and estuarine environments already experience much larger variations in CO_2_ levels (Sunda and Cai, 2012; Wallace et al., 2014), which will likely be exacerbated in the future (Cai et al., 2011; Melzner et al., 2013). These fluctuations may occur rapidly over minutes to hours, and reach levels as high as ~1 kPa (1% CO_2_, 10,000 μatm) (Borges *et al.*, 2006; Baumann *et al.*, 2015). This type of environmental hypercapnia implies a different physiological challenge compared to OA. Firstly, as environmental CO_2_ levels increase above CO_2_ levels in venous blood (typically ~0.3 kPa), CO_2_ diffusion gradients are reversed resulting in net uptake from the environment into the blood and inducing a much more pronounced respiratory acidosis (Tresguerres and Hamilton, 2017). Secondly, the sudden and extreme CO_2_ fluctuations must be met by equally fast, robust, and reversible acid-base regulatory responses.

Fish have a great capacity to restore blood pH to compensate for CO_2_-induced respiratory acidosis, which is largely achieved by excreting excess H^+^ and absorbing HCO_3_− using their gills (Brauner et al., 2019; Claiborne et al., 2002; Esbaugh, 2017; Evans et al., 2005; Perry and Gilmour, 2006). At the cellular level, these processes take place in specialized ion-transporting cells, or ionocytes. However, the underlying ion-transporting proteins and regulatory mechanisms are intrinsically different between freshwater and marine fishes and may also vary between species (Brauner and Baker, 2009; Claiborne et al., 2002; Evans et al., 2005; Perry and Gilmour, 2006). The few studies that have investigated acid-base regulation after acute exposure to ~1 kPa CO_2_ in marine fish have reported large variation of responses, with full blood pH compensation occurring between ~2 and 24 hours depending on the species (Hayashi et al., 2004; Larsen et al., 1997; Perry, 1982; Toews et al., 1983). Given the exquisite sensitivity of most proteins to changes in pH, variation in the time course of acid-base regulatory responses between species has important implications for whole organism performance. Haemoglobins (Hb) of fish species show strong Bohr and Root effects (Wells, 2009) which reduces Hb-O_2_ binding affinity and the O_2_ carrying capacity when erythrocyte intracellular pH (pH_i_) decreases. While fish have adaptations to regulate pH_i_ of erythrocytes (Cossins and Richardson, 1985; Nikinmaa and Tufts, 1989; Thomas and Perry, 1992), erythrocyte pH_i_ in many fish species (particularly marine fish) is closely linked to whole blood pH (Brauner and Baker, 2009; Shartau et al., 2020). Therefore, adaptations which enhance the speed of whole blood acid-base regulation will also provide faster restoration of O_2_ transport capacity and minimise disruption to energetically expensive activities such as foraging and digestion. However, little is known about why some species are able to compensate respiratory acidosis faster than others.

The gill ionocytes of marine fish excrete H^+^ using apical Na^+^-H^+^ exchangers (NHEs) driven by basolateral Na^+^/K^+^-ATPase (NKAs) (Brauner et al., 2019; Claiborne et al., 2002; Evans et al., 2005). Theoretically, H^+^ excretion during sudden exposure to hypercapnia could be upregulated by increased biosynthesis of NKA and NHE; however, transcriptional and translational responses typically take at least a few hours (e.g. Tresguerres *et al.*, 2005, 2006). Furthermore, protein turnover is energetically expensive (Pan et al., 2015), so short-term regulation of H^+^ excretion by synthesis and degradation of ion-transporting proteins would not be particularly efficient. Alternatively, the rapid upregulation of H^+^ excretion may be mediated by post-translational regulatory modifications such as insertion of pre-existing proteins into the ionocyte membrane, or morphological adjustments of its apical area, as reported by a variety of fishes in response to other acid-base disturbances (reviewed in Brauner et al., 2019; Tresguerres et al., 2019).

In the present study we investigated acid-base regulation of European sea bass, *Dicentrarchus labrax*, an active predator which seasonally inhabits shallow coastal, estuarine and saltmarsh environments (Doyle et al., 2017) where rapid and large fluctuations in CO_2_ levels occur (Borges et al., 2006). Specifically, we characterised blood acid-base regulation, erythrocyte intracellular pH (pH_i_) and O_2_ affinity, effects of seawater chemistry on speed of acid-base regulation, and changes in ionocyte NKA and NHE3 abundance, intracellular localisation, and apical surface area during acute exposure to ~1 kPa CO_2_.

## Methods

### Capture and Pre-Experimental Condition

Juvenile European sea bass were obtained by seine netting in estuaries and salt marshes from Dorset and the Isle of Wight on the south coast of the United Kingdom. Sea bass were transferred to the Aquatic Resources Centre of the University of Exeter where they were held in ~500 L tanks in a recirculating aquaculture system (RAS, total volume ~ 2500 L) at temperatures between 14 and 22°C. Sea bass were fed three times a week with commercial pellet (Horizon 80, Skretting) with a supplement of chopped frozen mussel (*Mytilus* edulis) once a week. For ~6 months before experiments sea bass were maintained at a temperature of 14°C and seawater CO_2_ levels of ~0.05-0.06 kPa (pH~8.10). Prior to all experimental procedures, sea bass were withheld food for a minimum of 72 hours. Animal collections were conducted under appropriate permits (Marine Management Organisation permit #030/17 & Natural England permit #OLD1002654) and all experimental procedures were carried out under home office licence P88687E07 and approved by the University of Exeter’s Animal Welfare and Ethical Review Board.

### Hypercapnia exposure

Individual sea bass were moved to isolation tanks (~20 L) and left to acclimate overnight for a minimum of 12 hours before exposure to hypercapnia. During the acclimation period isolation tanks were fed by the RAS at a rate of ~4 L min^−1^; with overflowing water recirculated back to the RAS. After overnight acclimation, hypercapnia exposure began by switching inflow of water from low CO_2_ control conditions to high CO_2_ seawater delivered from a header tank (~150 L) in which *p*CO_2_ levels had already been increased to ~1 kPa using an Aqua Medic pH computer (AB Aqua Medic GmbH). The pH computer maintained header tank *p*CO_2_ levels using an electronic solenoid valve which fed CO_2_ to a diffuser in the header tank if pH rose above 6.92 and stopped CO_2_ flow if pH dropped below 6.88. Additionally, to reduce CO_2_ fluctuation in isolation tanks during exposures, the gas aerating each tank was switched from ambient air to a gas mix of 1% CO_2_, 21% O_2_ and 78% N_2_ (G400 Gas mixing system, Qubit Biology Inc.). During exposures overflowing water from each isolation tank recirculated to the header tank creating an isolated experimental system of ~250 L. The experimental system was maintained at 14°C using a heater/chiller unit (Grant TX150 R2, Grant Instruments) attached to a temperature exchange coil in the header tank. To characterise the time course of acid-base regulation sea bass were exposed to hypercapnia (~1 kPa CO_2_) for either ~10 minutes, ~40 minutes, or ~135 minutes before measurements were taken. pH of isolation tanks was monitored with a separate pH probe and matched the header tank ~5 minutes after initial exposure. Measurements of an additional group of sea bass were obtained at normocapnic CO_2_ levels (~0.05 kPa CO_2_) to act as a pre-exposure control (hereafter this group is referred to as time = 0). At the time of sampling measurements of seawater pH (NBS scale), temperature and salinity, as well as samples of seawater to measure total CO_2_ (TCO_2_)/Dissolved Inorganic Carbon (DIC), were taken from each isolation tank. DIC analysis was conducted using a custom built system described in detail by Lewis *et al.* (2013). Measurements of pH, salinity, temperature and DIC were then input into CO2SYS (Pierrot et al., 2006) to calculate *p*CO2 and total alkalinity (TA) based on the equilibration constants refitted by Dickson and Millero (1987).

### Blood sampling and analysis

Following hypercapnia exposures (Table 1), sea bass were individually anaesthetised in-situ with a dose of 100 mg L^−1^ benzocaine. Blood samples were then obtained following the methodology outlined by Montgomery *et al.* (2019). The gill irrigation tank used was filled with water from the header tank and maintained at an appropriate CO_2_ level by aeration with the same gas mix feeding the isolation tanks. The water chemistry of the gill irrigation chamber was measured following the same procedures outlined for the isolation chambers, with one DIC sample taken at the end of blood sampling (Table S1).

**Table 1:** Mean ± s.e.m. of water chemistry parameters within isolation tanks during hypercapnia exposures prior to blood sampling.

|  | Duration |  |  |  |
| --- | --- | --- | --- | --- |
|  | 0 min<br>(Control) | ~10 min<br>(10.8 ± 0.26 min) | ~40 min<br>(41.0 ± 2.82 min) | ~135 min<br>(133.9 ± 2.27 min) |
| **Temperature (°C)** | 13.94 ± 0.04 | 13.90 ± 0.03 | 13.94 ± 0.02 | 13.89 ± 0.04 |
| **pH (NBS)** | 8.15 ± 0.01 | 6.98 ± 0.01 | 6.96 ± 0.01 | 6.95 ± 0.01 |
| **Salinity** | 34.1 ± 0.1 | 34.1 ± 0.1 | 34.7 ± 0.2 | 34.0 ± 0.1 |
| ***p*CO<sub>2</sub> (kPa)** | 0.059 ± 0.001 | 0.898 ± 0.022 | 0.944 ± 0.072 | 0.945 ± 0.017 |
| **TA (µM)** | 3,251 ± 20 | 2,804 ± 16 | 2,833 ± 11 | 2,777 ± 29 |

Immediately after sampling, extracellular pH was measured on 30 μL of whole blood using an Accumet micro pH electrode and Hanna HI8314 pH meter calibrated to 14 °C using pH_NBS_ 7.04 and 9.21 appropriate buffers. Measurements of blood pH were made in a temperature-controlled water bath. Three 75 μL micro capillary tubes were then filled with whole blood and anaerobically sealed with Critoseal capillary tube sealant (Fisher) and paraffin oil and centrifuged for 2 minutes at 10,000 rpm. Haematocrit (Hct) was measured using a Hawksley micro-haematocrit reader. Plasma was then extracted from capillary tubes for analysis of TCO_2_ using a Mettler Toledo 965D carbon dioxide analyser. Plasma *p*CO_2_ and HCO_3_− were then calculated from TCO_2_, temperature and blood pH using the Henderson-Hasselbalch equation with values for solubility and pK^1^_app_ based on Boutilier *et al.* (1984, 1985). Haemoglobin (Hb) content of 10 μL of whole blood was also assessed by the cyanmethemoglobin method (after addition to 2.5 mL of Drabkin’s reagent, Sigma). Half the remaining whole blood was centrifuged at 10,000 rpm for 2 minutes at 4°C. The resulting plasma was separated and 10 μL was diluted in ultrapure water, snap frozen in liquid N_2_, and stored at −80°C before later being used to measure plasma cation and anion concentrations using ion chromatography (Dionex ICS 1000 & 1100, Thermo-Scientific, UK). The remaining plasma was snap frozen in liquid N_2_ and stored at −80°C before measurements of plasma lactate and glucose were made (YSI 2900D Biochemistry Analyzer, Xylem Analytics). After separating the plasma, the surface of the leftover erythrocyte pellet was blotted to remove the leukocyte layer. The erythrocyte pellet was then snap frozen in liquid nitrogen for 10 seconds and thawed in a 37°C water bath for 1 minute prior to intracellular pH (pH_i_) measurements as described by Zeidler and Kim (1977), and validated by Baker *et al.* (2009). All measurements or storage of blood occurred within 10 minutes of blood sampling. Finally, Hb-O_2_ affinity was measured following the methods outlined in Montgomery *et* al. (2019) using a Blood Oxygen Binding System (BOBS, Loligo systems), as detailed by Oellermann *et al.* (2014).

### Flux measurements

The flux of acid-base relevant ions between sea bass and seawater was measured over a ~135 minute time period in normocapnic conditions (n = 7) and immediately following exposure to hypercapnia (n = 8, Table 2). At the start of the measurements the flow to the isolation tanks was stopped and water chemistry maintained at the desired *p*CO_2_ by gassing the tanks with either ambient air (control) or a 1% CO_2_ gas mix (hypercapnia). Seawater samples for measuring TA were taken at the beginning and end of the ~135 minute flux period, preserved by adding 40 μL of 4% (w/v) mercuric chloride per 10 mL of seawater, and stored at 4 °C (Dickson et al., 2007) prior to analysis by double titration using an autotitrator (Metrohm 907 Titrando with 815 Robotic USB Sample Processor XL, Metrohm). TA measurements were made using a double titration method modified from Cooper *et al.* (2010) as detailed by Middlemiss *et al.* (2016). Briefly, 20 mL samples were titrated to pH 3.89 using 0.02 M HCl whilst gassing with CO_2_-free N_2_, pH was then returned to starting values by titrating with 0.02 M NaOH. Samples for measuring total ammonia were frozen at −20°C before ammonia concentration was measured using a modified version of the colourimetric method of Verdouw *et al.* (1978) at 660 nm using a microplate reader (NanoQuant infinite M200 pro, Tecan Life Sciences). A calibration curve was constructed using NH_4_Cl standards in seawater.

**Table 2:** Mean ± s.e.m. of water chemistry parameters within individual tanks during flux measurements.

| Treatment | Duration (min) | pH (NBS) | Temperature (°C) | Salinity | $p\text{CO}_2$ (kPa) | TA ($\mu\text{M}$) |
| --- | --- | --- | --- | --- | --- | --- |
| Normocapnia | 132.3 ± 0.9 | 7.96 ± 0.01 | 14.03 ± 0.04 | 33.6 ± 0.2 | 0.072 ± 0.002 | 2395 ± 21 |
| Hypercapnia | 123.6 ± 2.4 | 6.91 ± 0.03 | 13.90 ± 0.12 | 33.6 ± 0.0 | 0.862 ± 0.056 | 2299 ± 33 |

Acid-base relevant fluxes (μmol kg^−1^ h^−1^) were then calculated using the following equation:

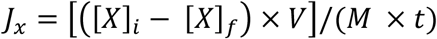

as described by Wilson and Grosell (2003), where V is the volume of water (L) in the isolation tank (after the initial sample is taken), M is the mass of the sea bass (kg), t is the duration of the flux period (h) and [X]_i_ and [X]_f_ are the ion concentrations in the chamber water (μmol L^−1^) at the beginning and end of the flux period. By reversing the initial and final values titratable acid, instead of base, fluxes can be calculated so that a positive value equals acid uptake (i.e. HCO_3_− excretion) and a negative value equals acid excretion (i.e. HCO_3_− uptake). We then calculated net acid-base fluxes (μeq kg^−1^ h^−1^) as the sum of titratable acid and total ammonia (T_amm_) flux (McDonald and Wood, 1981).

### Exposure to Low Total Alkalinity

Sufficient 1 M HCl was added to ~250 L seawater to reduce TA by over 90% from ~2,800 μM to ~200 μM, followed by overnight aeration to equilibrate CO_2_ with atmospheric levels. We then adjusted the *p*CO_2_ of the low TA seawater to the desired level of ~1 kPa as described above, and a pH set point of 5.75. Sea bass were placed in the individual isolation boxes (fed by the RAS as detailed for normal TA hypercapnia exposures) and left to acclimate overnight before being exposed to the combined low TA and hypercapnia treatment. Flow to individual isolation boxes was stopped, and ~75% of the water from the isolation box was drained and refilled with low TA, hypercapnic water. This process was repeated 3 times over a period of ~5 minutes. The gas mix aerating each isolation box was switched from ambient air to a 1% CO_2_ gas mix to maintain the desired *p*CO_2_ levels. After ~135 minutes exposure, each seabass was anaesthetised and sampled for blood acid-base measurements as detailed previously. The water chemistry of isolation boxes (Table S2) and gill irrigation chambers (Table S3) was measured at the time of blood sampling.

### Gill sampling

Gill tissue was sampled from sea bass exposed to ambient CO_2_ conditions (n = 5) and to hypercapnia for ~135 minutes (n=5, taken immediately after the flux measurements) in normal TA seawater (Table S4). Mean water chemistry conditions during flux measurements (Table 2) and experienced by sea bass prior to gill sampling (Table S4) differ because gill samples were only collected from 5 of the 8 sea bass used for flux measurements. After euthanizing the sea bass in an anaesthetic bath (benzocaine, 1 g L^−1^), gills were dissected and rinsed in phosphate buffered saline (PBS). The first gill arch on the left side was flash frozen in liquid N_2_ and stored at −80°C for Western blotting, and the first gill arch on the right side was fixed in 4% paraformaldehyde in 0.1 M phosphate buffer saline (PBS) (diluted from 16% electron microscopy grade paraformaldehyde. Electron Microscope Science catalogue # 15710), overnight at 4°C for immunohistochemistry. Following a ~10-hour fixation, gill samples were transferred to 50% ethanol for ~10 hours at 4 °C, and then stored in 70% ethanol at 4°C.

### Antibodies

NKA was immunodetected using α5, a mouse monoclonal antibody against the α-subunit of chicken NKA (a5, Developmental Studies Hybridoma Bank, Iowa City, IA, USA; Lebovitz *et al.*, 1989). This antibody universally recognizes NKA in teleost fishes including yellowfin tuna (*Thunnus albacares*; Kwan *et al.*, 2019), Pacific chub mackerel (*Scomber japonicus*; Kwan *et al.*, 2020), and California killifish (*Fundulus parvipinnis*; Nadler *et al.*, 2021). Rabbit anti-NHE3 polyclonal antibodies were generously donated by Dr Junya Hiroi (St. Marianna University School of Medicine, Kawaski, Japan); they target two epitope regions within rainbow trout (*Oncorhynchus mykiss*) NHE3b (GDEDFEFSEGDSASG and PSQRAQLRLPWTPSNLRRLAPL), and recognize NHE3 of multiple teleost species including European sea bass (*D. labrax*; Blondeau-Bidet *et al.*, 2019). The secondary antibodies were goat anti-mouse HRP-linked and goat anti-rabbit HRP-linked (Bio-Rad, Hercules, CA, USA) for immunoblotting, and goat anti-mouse Alexa Fluor 546 and goat anti-rabbit Alexa Fluor 488 (Invitrogen, Grand Island, USA) for immunohistochemistry.

### Western Blotting

Western blotting followed the procedures outlined in Kwan *et al.*, (2019, 2020). While frozen on dry ice, the gill filament and lamellae were separated from the gill arch using a razor blade. The excised tissue was then immersed in liquid N_2_ and pulverized in a porcelain grinder, then submerged within an ice-cold, protease inhibiting buffer (250 mmol L^−1^ sucrose, 1 mmol L^−1^ EDTA, 30 mmol L^−1^ Tris, 10 mmol L^−1^ benzamidine hydrochloride hydrate, 1 mmol L^−1^ phenylmethanesulfonyl fluoride, 1 mmol L^−1^ dithiothreitol, pH 7.5). Samples were further homogenized using a handheld VWR Pellet Mixer (VWR, Radnor, PA, USA) for 15 second intervals (3 times) while on ice. Next, samples were centrifuged (3,000 g, 4°C; 10 minutes), and the resulting supernatant was considered the crude homogenate. An aliquot of the crude homogenate was further subjected to a higher speed centrifugation (21,130xg, 4°C; 30 minutes), and the pellet was saved as the membrane-enriched fraction. Bradford assay was used to determine protein concentration (Bradford, 1976), which was used to normalize protein loading.

On the day of Western blotting, samples were mixed with an equal volume of 90% 2x Laemmli buffer and 10% β-mercaptoethanol. Samples were then heated at 70°C for 5 minutes, and the proteins (20 μg per lane) were loaded onto a polyacrylamide mini gel (4% stacking; 10% separating) – alternating between control and high CO_2_ treatments to avoid possible gel lane effects. The gel ran at 60 volts for 15 minutes, then 100 V for 1.5 hours, and proteins were transferred onto a polyvinylidene difluoride (PVDF) membrane using a wet transfer cell (Bio-Rad) at 70 volts for 2 hours at 4 °C. PVDF membranes were then incubated in Tris-buffered saline with 1% tween (TBS-T) with milk powder (0.1 g/mL) at room temperature for 1 hour, then incubated with primary antibody (NKA: 10.5 ng/ml; NHE3: 1:1,000) in blocking buffer at 4°C overnight. On the following day, PVDF membranes were washed in TBS-T (three times; 10 minutes each), incubated in blocking buffer with secondary antibodies (1:10,000) at room temperature for 1 hour, and washed again in TBS-T (three times; 10 minutes each). Bands were made visible through addition of ECL Prime Western Blotting Detection Reagent (GE Healthcare, Waukesha, WI) and imaged with the Universal III Hood (BioRad). Following imaging, the PVDF membrane was incubated in Ponceau stain (10 minutes, room temperature) to estimate protein loading. Relative NKA and NHE protein abundance (n = 5 per treatment) was quantified using the Image Lab software (version 6.0.1; BioRad) and normalized by the protein content in each lane.

### Immunohistochemistry

Immunohistochemistry was performed as described in Kwan *et al.*, (2020). Fixed gill tissue stored in 70% ethanol was rehydrated in PBS + 0.1% tween (PBS-T) for 10 minutes, and gill filaments were dissected out to ease subsequent imaging. Autofluorescence was quenched by rinsing in ice-cold PBS with sodium borohydride (1.0 – 1.5 mg mL^−1^; six times; 10 minutes each), followed by incubation in blocking buffer (PBS-T, 0.02% normal goat serum, 0.0002% keyhole limpet haemocyanin) at room temperature for one hour. Samples were incubated with blocking buffer containing primary antibodies (NKA: 40 ng/mL; NHE3: 1:500 [c.f. Seo *et al.*, (2013)]) at 4°C overnight. On the following day, samples were washed in PBS-T (three times at room temperature; 10 min each), and incubated with the fluorescent secondary antibodies (1:500) counterstained with DAPI (1 μg mL^−1^) at room temperature for 1 hour. Samples were washed again in PBS-T as before, then placed on a concave slide for imaging using an inverted confocal microscope (Zeiss LSM 800 with Zeiss ZEN 2.6 blue edition software; Cambridge, United Kingdom). Samples incubated without primary antibodies had no signal (Fig. S1).

### Quantification of Ionocyte Apical Surface Area

The apical surface area of gill ionocytes were quantified through a combination of whole-mount imaging (40X objective lens with deionized water immersion), optical sectioning, and XZ- and YZ-plane analysis. A relatively flat surface on the gill filament was selected under 0.5x scanning confocal magnification to ensure imaging would be performed on ionocytes in an upright position thus minimizing errors in apical surface area quantification due to angle distortion. After locating an ionocyte by its distinctive NKA signal, the scanning confocal magnification was increased to 5.0X and the entire cell was Z-stack imaged (optimal interval automatically selected: 0.07-0.12 μm per slice). Subsequent viewing of the Z-stack from the X-Z and Y-Z planes allowed us to assess intracellular localisation, and to identify the image slice that captured the entire apical surface (typically, the second slice from the top of the cell). Next, the ionocyte’s apical surface area (identified by NHE3 immunofluorescence signal) was quantified using FIJI (Schindelin et al., 2012). For each sea bass (n=5 per treatment), the average apical surface area was calculated from three ionocytes from different gill filaments.

### Statistical Analysis

All statistical analysis was performed using R v3.6.3 (R Core Team, 2020). Changes in blood chemistry parameters over time in response to hypercapnia exposure were analysed using one-way ANOVA before assumptions of equal variances of data and normality of model residuals were checked. Post-hoc tests were conducted on least-square means generated by package ‘emmeans’ (Lenth, 2020), with Tukey adjusted p-values for multiple comparisons. Some data did not meet required assumptions for one-way ANOVA. Unequal variances were observed in measurements of *p*CO_2_ between treatments, as such we used Welch’s ANOVA with Tukey’s pairwise comparisons using Benjamini-Hochberg corrections for post-hoc testing. Measurements of blood pH and P_50_ did not meet assumptions of normality and were analysed using the Kruskal-Wallis test with post-hoc comparisons made with Dunn’s test from package ‘FSA’ (Ogle et al., 2020), using Benjamini-Hochberg corrections for multiple comparisons. As a result of unusually high measurements of plasma [Cl^−^] and [Na^+^] in some samples a ROUT test was conducted (Q = 0.5%) in Graphpad Prism 9 to identify potential outliers (Motulsky and Brown, 2006). Samples in which plasma [Cl^−^] and [Na^+^] were both identified as outliers by the ROUT test were excluded from the dataset prior to subsequent statistical analysis. Flux measurements were analysed using Student’s t-test after checking data met assumptions of normality and equal variance. Relative protein abundance and ionocyte apical area met both assumptions of normality and equal variance and were analysed using one-tailed t-test (control response < CO_2_-exposed response).

## Results

### Blood chemistry

Exposure to environmental hypercapnia caused significant changes in blood pH of sea bass over time (Kruskal-Wallis test, χ^2^ = 25.0, df = 3, p < 0.001). There was a pronounced acidosis of the blood from pH 7.84 (± 0.02) in control conditions (normocapnia, time = 0) to 7.50 (± 0.03) after exposure to hypercapnia for ~10 minutes (Fig. 1A, D). Following this initial acidosis sea bass completely restored blood pH to control levels after ~135 minutes (Fig. 1A, D). Blood pH regulation was accompanied by a ~5-fold increase in plasma HCO_3_−, from 4.5 ± 0.3 to 21.9 ± 0.7 mM, over the ~135 minute exposure (Fig. 1C, D, One-way ANOVA, F = 203.3, df = 3, p < 0.001).

**Figure 1:**
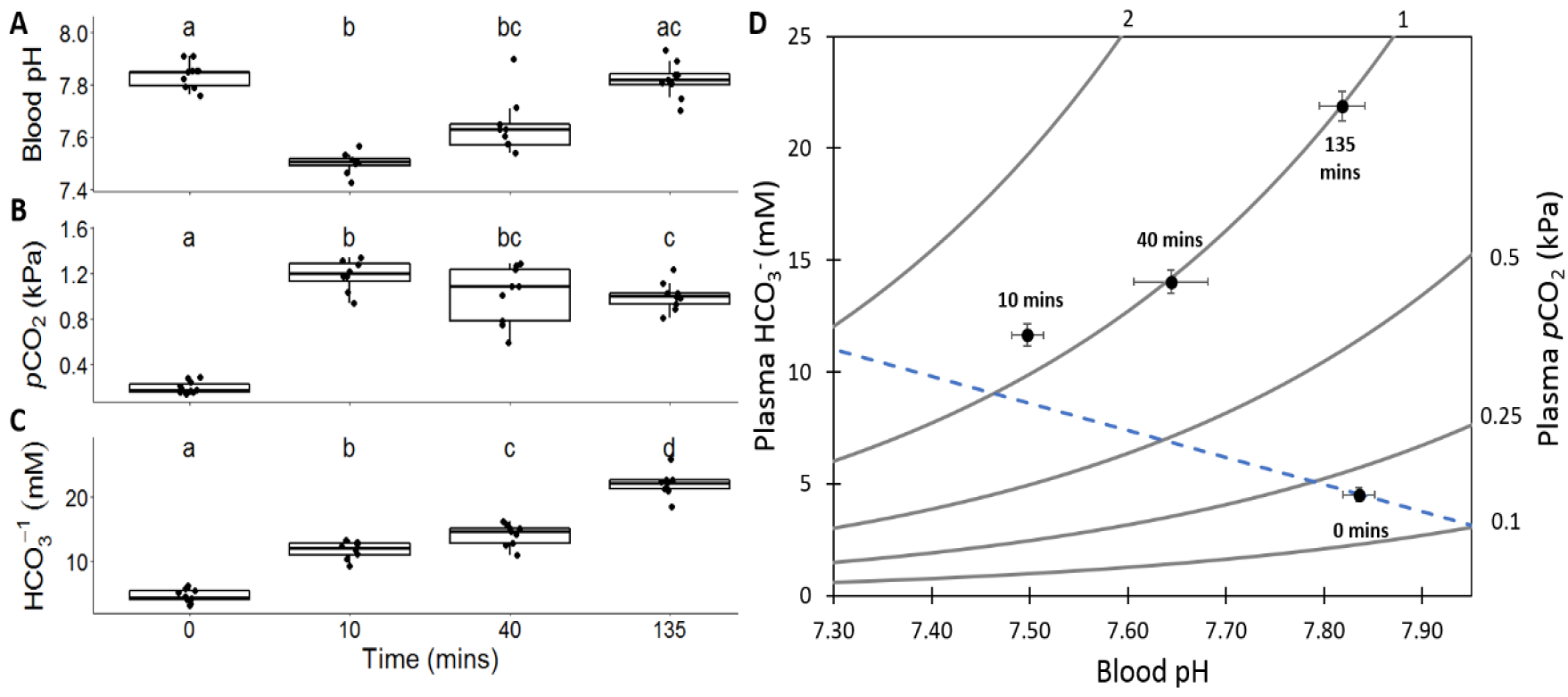
Changes in **A.** blood pH, **B.** plasma *p*CO_2_, and **C.** plasma HCO_3_− between European sea bass in control conditions (~0.05 kPa CO_2_, Time = 0, n= 10) and after exposure to ~0.9 kPa CO_2_ for ~10 minutes (n = 8), ~40 minutes (n = 9), and ~135 minutes (n = 9). Significant differences between parameters at each time point are indicated by different lower case letters (**A.** Dunn’s test, p < 0.05; **B.** Pairwise comparison using Benjamini-Hochberg correction, P < 0.05 **C.** Pairwise comparison of least square means, p < 0.05) **D.** Combined changes of all three acid-base parameters are expressed as a pH/HCO_3_−/*p*CO_2_ diagram (blue dashed line indicates estimated non-bicarbonate blood buffer line based on equations from Wood *et al.* (1982)) values represent mean ± s.e.m.

Plasma *p*CO_2_ showed significant changes during exposure to hypercapnia (Fig. 1B, D, Welch’s ANOVA, F = 202.5, df = 3, p < 0.001). The initial decrease in blood pH of sea bass was driven by a rapid and large (~6-fold) increase in plasma *p*CO_2_, from 0.200 ± 0.016 to 1.185 ± 0.049 kPa CO_2_, within the first 10 minutes of exposure. There was a small but significant decline in plasma *p*CO_2_ between sea bass sampled ~10 minutes after exposure and sea bass sampled ~135 minutes after exposure (Fig. 1B). There were no significant differences in plasma glucose or lactate levels between any treatment groups with values for all sea bass of 5.90 ± 0.43 mM and 0.45 ± 0.05 mM respectively.

### Flux measurements

Sea bass switched from slight net base excretion under control normocapnic conditions to net acid excretion that was ~2.5-fold larger in magnitude during 135 minutes of hypercapnia (Fig. 2C, Student’s t-test, t = −2.25, df = 13, p = 0.042). This was driven by a switch from a small apparent HCO_3_− excretion to a large apparent HCO_3_− uptake (Fig. 2A). There were no significant differences in T_Amm_ excretion (Fig. 2B).

**Figure 2:**
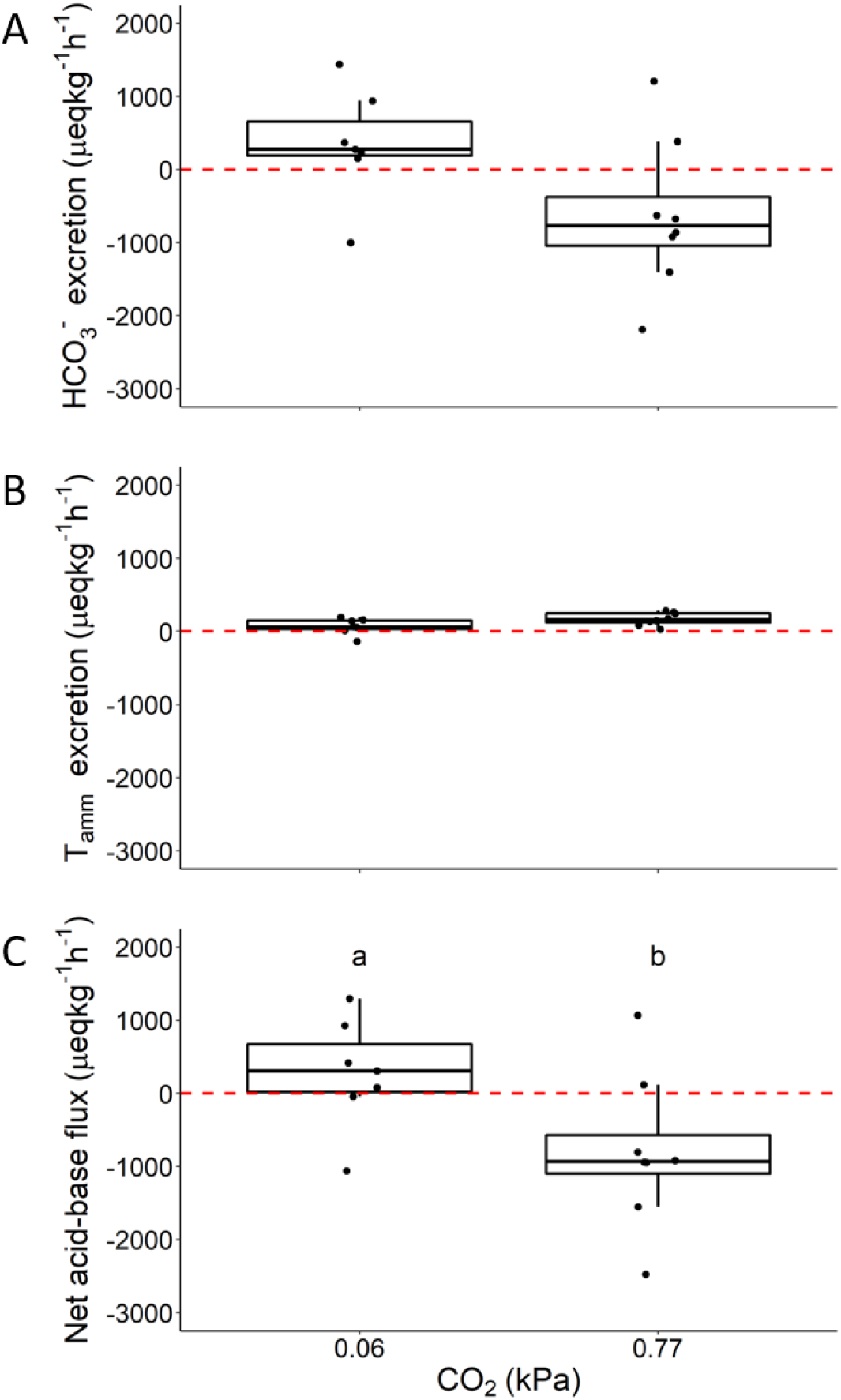
Changes in **A.** excretion of HCO_3-,_ **B.** excretion of total ammonia (T_amm_) and **C.** net acid-base flux between European sea bass in control conditions (n = 7, ~0.07 kPa CO_2_) and after ~135 minutes exposure to hypercapnia (n = 8, ~0.84 kPa CO_2_). Significant difference in parameters are indicated by different lower-case letters (Student’s t-test, p < 0.05).

### Oxygen transport capacity

The initial drop in blood pH during hypercapnia exposure was reflected in changes in erythrocyte pH_i_ (Fig. 3A, One-way ANOVA, F = 12.34, df = 3, p < 0.001) and erythrocyte [H^+^] (Fig. 3B, One-way ANOVA, F = 14.64, df = 3, P < 0.001). However, erythrocyte pH_i_ and [H+] returned to control levels after ~40 minutes of exposure to hypercapnia (Fig. 3A, B). As expected, the significant changes in erythrocyte pH_i_ and [H+] affected haemoglobin-O_2_ binding affinity leading to a ~3-fold increase in P_50_ after 10 minutes, from 1.78 kPa O_2_ (± 0.30 kPa O_2_) in control sea bass to 5.60 kPa O_2_ (± 0.36 kPa O_2_) (Kruskall-wallis test, χ^2^ = 17.4, df = 3, p < 0.001). There were no changes in Hills’ number across hypercapnia exposure (One-way ANOVA, F = 1.48, df = 3, p = 0.248). The rapid recovery of erythrocyte pH_i_ and [H+] after ~40 minutes led to P_50_ returning to pre-exposure levels (Fig. 3B).

**Figure 3:**
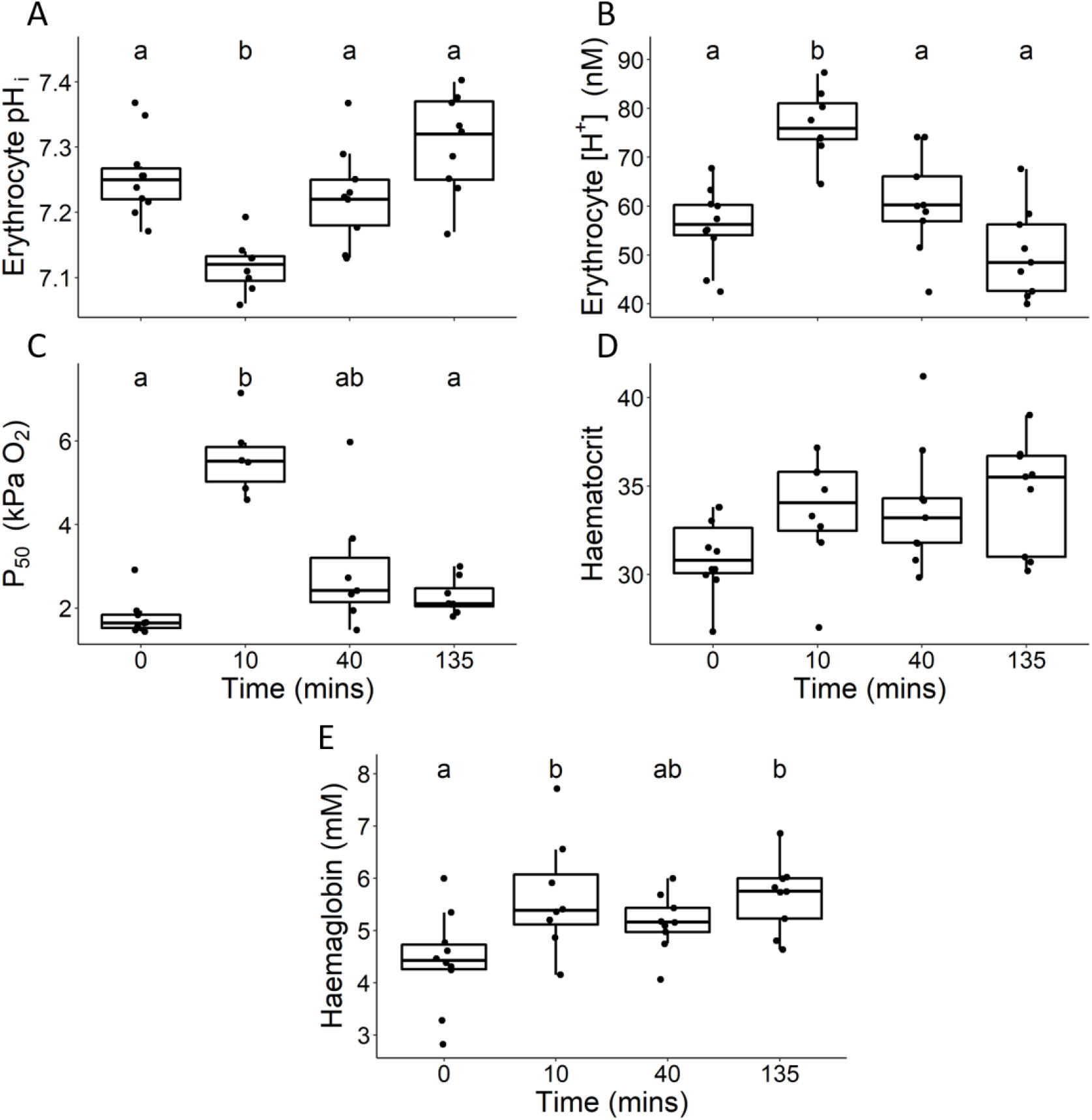
Changes in **A.** erythrocyte intracellular pH (erythrocyte pH_i_), **B.** erythrocyte [H^+^], **C.** haemoglobin-O_2_ binding affinity (P_50_), **D.** haematocrit and **E.** haemoglobin level between European sea bass in control conditions (~0.05 kPa CO_2_, Time = 0) and after exposure to ~0.9 kPa CO_2_ for ~10 minutes, ~40 minutes, and ~135 minutes. Significant differences between parameters at each time point are indicated by different lowercase letters (**A.**, **B., D.** and **E.** Pairwise comparisons of least square means, p < 0.05, **C.** Dunn’s test, p < 0.05).

Sea bass exposed to hypercapnia also experienced a ~25% increase in haemoglobin levels (Fig. 3D), at ~10 minutes and ~ 135 minutes compared to control sea bass (One-way ANOVA, F = 4.60, df = 3, p = 0.009). In addition, sea bass exposed to hypercapnia exhibited an ~8-10% increase in haematocrit (Fig. 3C), although this increase was marginally non-significant (One-way ANOVA, F = 2.40, df = 3, 0.086).

### Response to Hypercapnia in Seawater with Low Total Alkalinity

To test the influence of environmental availability of [HCO_3_−] on acid-base regulation, a group of sea bass were exposed to hypercapnia in low alkalinity seawater. These sea bass were unable to compensate for a respiratory acidosis when exposed to acute hypercapnia for ~135 minutes (Fig. 4A). Blood pH was 0.37 units (95 % CI = 0.33-0.41) lower than sea bass exposed to hypercapnia in normal alkalinity seawater for the same length of time and the same blood pH as we recorded in sea bass ~10 minutes after exposure to hypercapnia in normal seawater. Additionally, sea bass in low alkalinity seawater did not actively accumulate HCO_3_− when exposed to environmental hypercapnia for ~135 minutes (Fig. 4C). Indeed, the 11.8 mM increase in plasma [HCO_3_−] (95% CI = 10.6-12.9 mM) followed the predicted non-bicarbonate buffering line (Fig. 4D).

**Figure 4:**
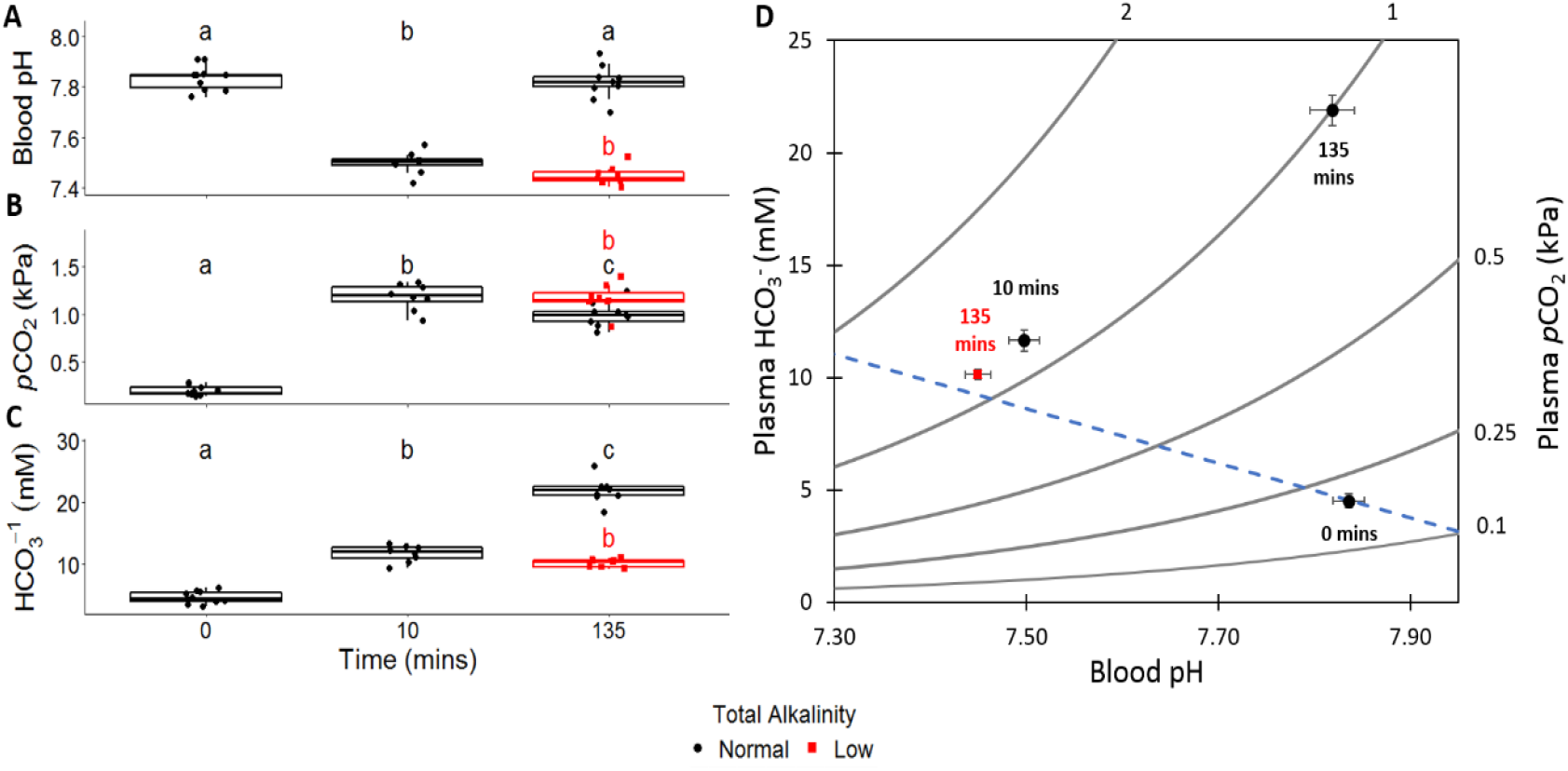
Comparison of **A.** blood pH, **B.** plasma *p*CO_2_ and **C.** plasma HCO_3_− between European sea bass in control conditions (n = 10, Time = 0), exposed to hypercapnia for ~10 minutes (n = 8) in normal (~2800 μM) total alkalinity (TA) seawater, exposed to hypercapnia for ~135 minutes in normal (~2800 μM) TA seawater (n = 9), and exposed to hypercapnia for ~135 minutes in low (~200 μM) TA seawater (n =8). Significant differences between parameters at each time point are indicated by different lower-case letters (Pairwise comparison of least squares means, p<0.05). For measurements taken after ~135 minutes of exposure to hypercapnia the colour indicates the TA treatment (i.e. black = normal TA, red = low TA). **D.** Combined changes of all three acid-base parameters are expressed as a pH/HCO_3_−/*p*CO_2_ diagram (blue dashed line indicates estimated non-bicarbonate blood buffer line based on equations from Wood *et al.* (1982)) values represent mean ± s.e.m.

### Plasma Ion Concentrations

Plasma [Cl^−^] significantly decreased by 13.1 mM (95% CI = 10.1-16.2 mM) from 134.9 mM (± 5.1) in sea bass in normocapnia to 121.7 mM (± 1.3) in sea bass exposed to hypercapnia for ~135 minutes (Kruskall-Wallis test, χ^2^ = 11.1, df = 4, p = 0.025). This decrease in plasma [Cl^−^] was not seen in sea bass exposed to hypercapnia in low alkalinity water (Fig. 5A). Decreases in plasma [Cl^−^] showed a correlation with increasing bicarbonate (Figure 5A inset, Kendall’s tau correlation, τ = −0.32, p = 0.005). Plasma [Na^+^] showed no significant changes over the time course of hypercapnia exposure (One-way ANOVA, F = 1.063, p = 0.391), and there were no differences in [Na^+^] after ~135 minutes of exposure to hypercapnia between sea bass in normal and low TA seawater (Fig. 5B). As such, there was no correlation between plasma [Na^+^] and [H^+^] (Fig. 5B inset, Kendall’s tau correlation, τ = 0.08, p = 0.471).

**Figure 5:**
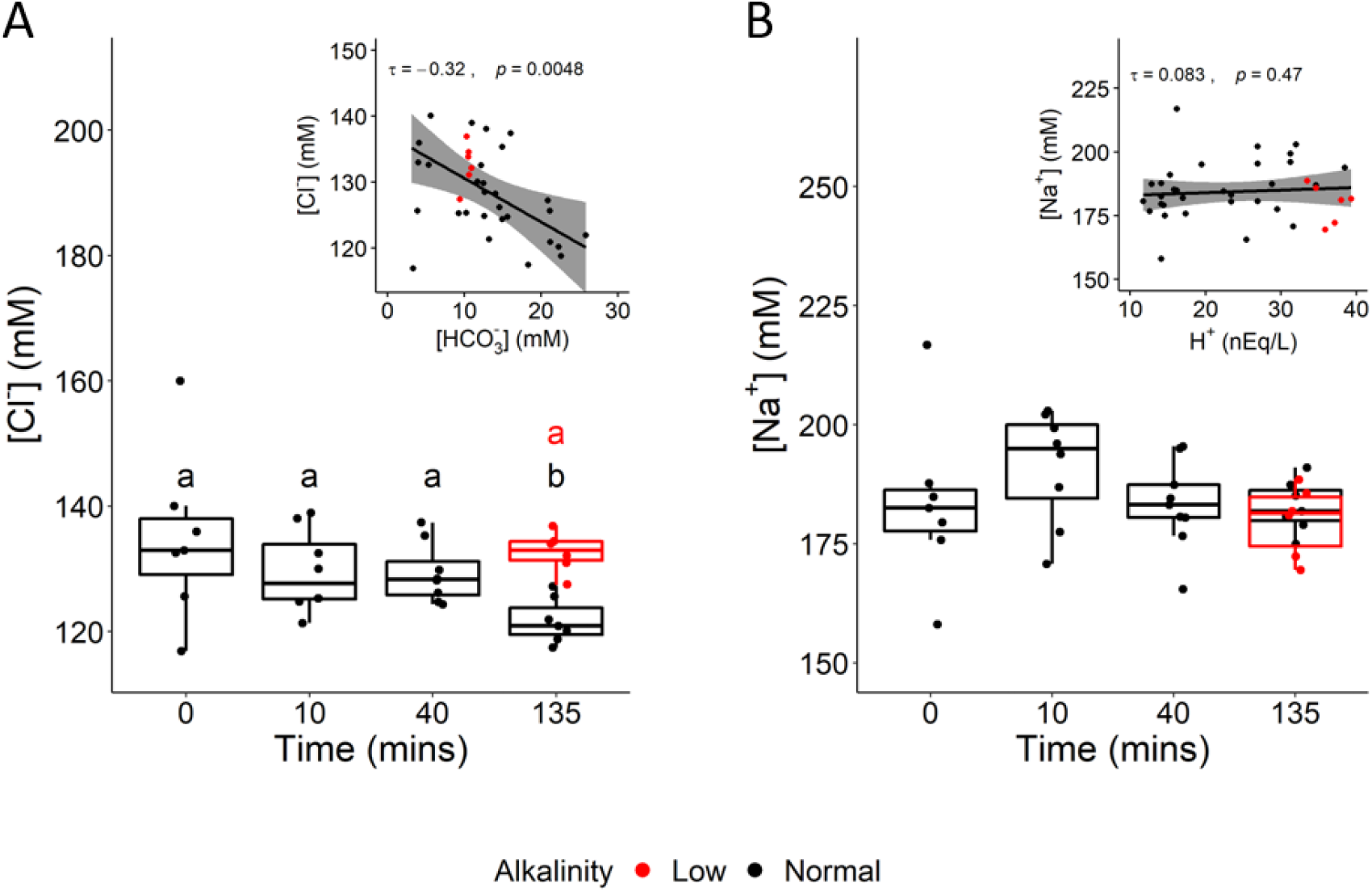
Comparison of **A.** plasma [Cl^−^] and **B.** plasma [Na^+^] between European sea bass in control conditions (n = 7, Time = 0), exposed to hypercapnia for ~10 minutes (n = 8), ~40 minutes in or ~135 minutes in normal (~2800 μM) TA seawater (n = 7), and exposed to hypercapnia in low (~200 μM) TA seawater (n = 6). Significant differences between [Cl^−^] at each time point are indicate by different lower-case letters (Pairwise comparison of least squares means, p<0.05). Insets show correlation between **A.** plasma [Cl-] and [HCO_3_−]. **B.** plasma [Na^+^] and [H^+^]. T and p value shown represent results of Kendall’s tau correlation. Shaded area represents 95% CI of linear regression between measures. For insets and measurements taken after ~135 minutes of exposure to hypercapnia the colour indicates the TA treatment (i.e. black = normal TA, red = low TA).

### NKA and NHE3 Protein Abundance

Exposure to hypercapnia did not induce significant changes in the abundance of NKA or NHE3 in crude homogenates (indicative of total protein abundance) or the abundance of NKA and NHE3 in membrane-enriched fractions (indicative of protein that was present in the apical or basolateral plasma membranes) (One-tailed t-test, P > 0.05; Fig. 6).

**Figure 6:**
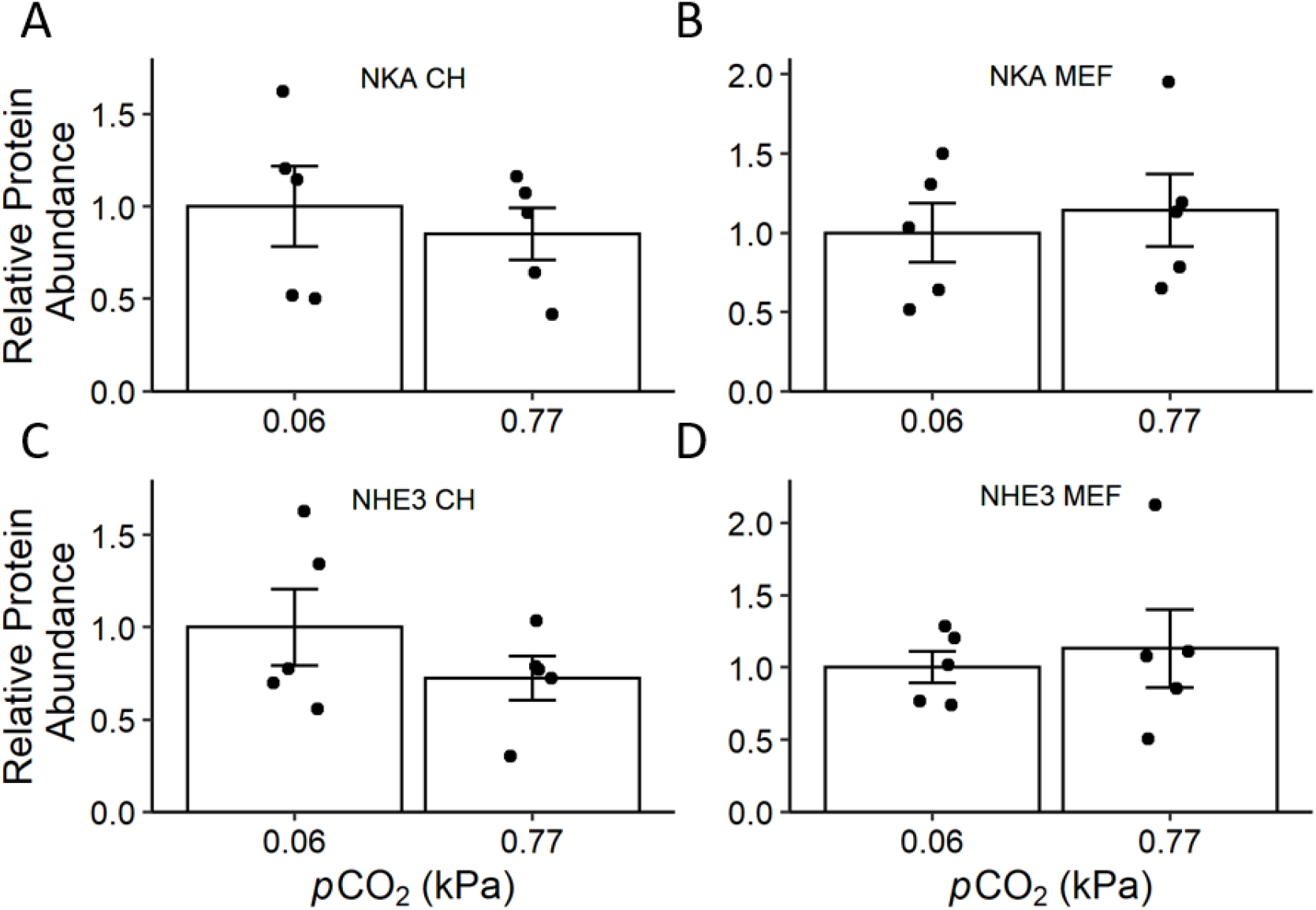
Comparison of gill **A.** Na^+^/K^+^-ATPase (NKA) in crude homogenates (CH), **B.** NKA in the membrane-enriched fraction (MEF) as well as **C.** Na^+^/H^+^ Exchanger 3 (NHE3) CH, and **D.** NHE3 MEF abundance between European sea bass exposed to control conditions (~0.06 kPa CO_2_) and to hypercapnia (~0.77 kPa CO_2_) for ~135 minutes (n = 5 per treatment, 1-tailed t-test). Bars show mean ± s.e.m., points show raw data; there were no significant differences between any measurements (1-tailed t-tests, p > 0.05).

### Ionocyte Intracellular Localisation and Apical Surface Area

NKA-rich ionocytes were primarily localised on the gill filament trailing edges and the basal portion of the gill lamellae (Fig. S1), all NKA-rich ionocytes also expressed NHE3 in their apical region (Fig. 7A). Despite analysis using high magnification imaging, optical sectioning, and XZ- and YZ-plane visualization, we found no evidence of NHE3 intracellular localisation (Figure 7B, B’, C, C’). The ionocyte’s apical surface area (based on the NHE3 signal) significantly increased, almost doubling from 3.38 ± 0.41 to 6.45 ± 0.64 μm^2^, after exposure to ~135 min of hypercapnia (one-tailed t-test, t = 4.048, df = 6.828, p = 0.003; Fig. 7D).

**Figure 7:**
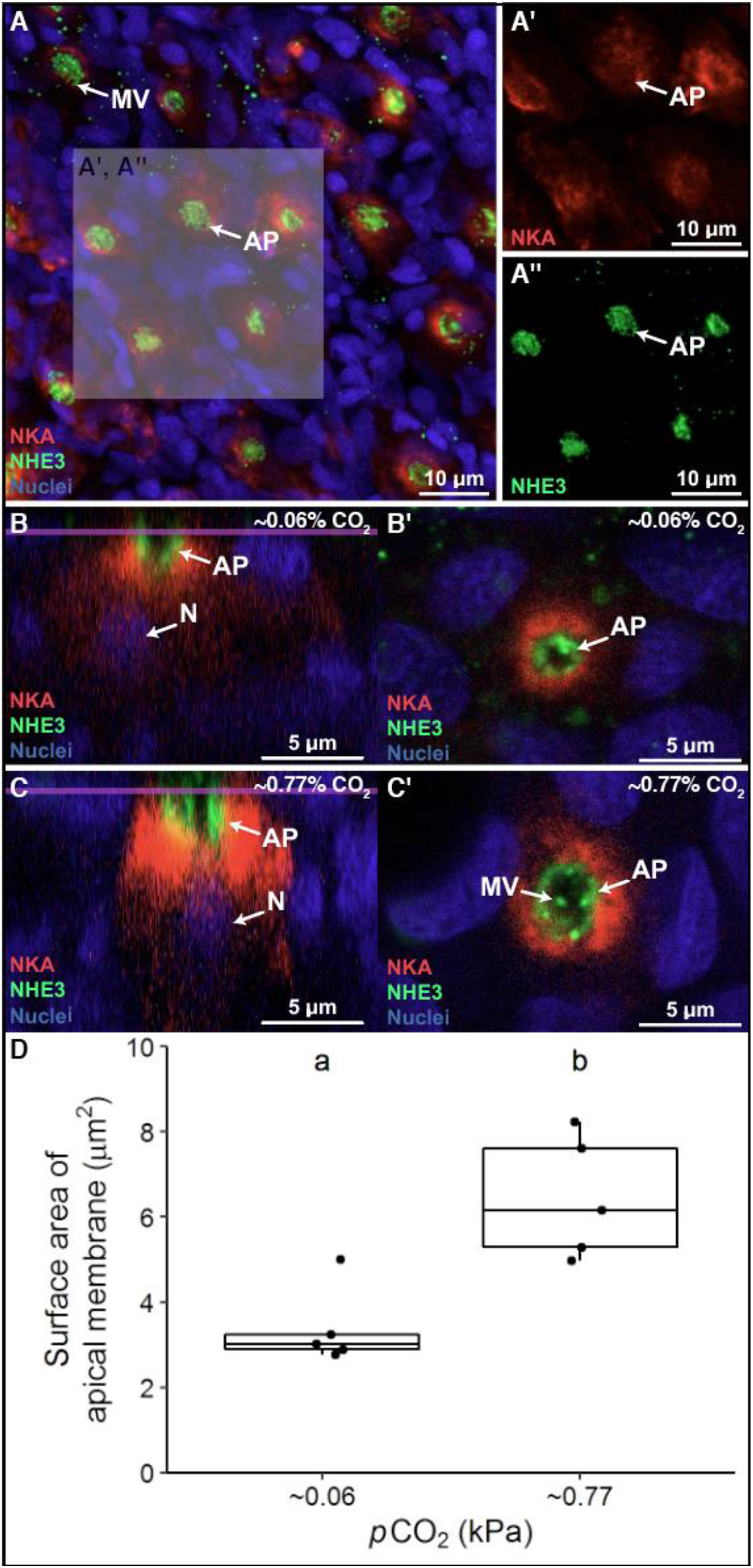
**A**. The European sea bass’ gill ionocytes express abundant **A’.** basolateral Na^+^/K^+^-ATPase (NKA, red) and **A’’.** apical Na^+^/H^+^ Exchanger 3 (NHE3, green). Comparison of gill ionocytes between European sea bass exposed to **B, B’.** control conditions (~0.06 kPa CO_2_) and **C, C’.** to ~0.77 kPa CO_2_ for ~135 minutes revealed no changes in intracellular localisation, but determined **D.** hypercapnia-exposed sea bass had significantly wider apical surface area (n = 5 per treatment, 1-tailed t-test, p = 0.003). The purple line in **B** and **C** denotes the slice at which **B’** and **C’** were imaged. Nuclei (blue) are stained with DAPI.

## Discussion

Our results indicate that European sea bass are able to rapidly compensate for hypercapnia-induced blood acidosis when the environmental CO_2_ is at the extreme high end of the spectrum encountered in their natural habitat. Complete restoration of blood pH after exposure to ~1 kPa CO_2_ was achieved within ~2 hours via a switch from net base excretion to net acid excretion and a subsequent accumulation of HCO_3_− in plasma. In addition, erythrocyte pH_i_ and Hb-O_2_ affinity were restored to pre-exposure levels after just ~40 min, and there was a 20% increase in the blood haemoglobin concentration together with a trend for ~10% haematocrit increase. These results suggest an adrenergic response that stimulates a β-NHEs in the erythrocytes (Nikinmaa, 2012), and contracts the spleen resulting in the release of erythrocytes into the circulation (Crocker and Cech, 1997; Lee et al., 2003; Perry and Kinkead, 1989; Vermette and Perry, 1988). The end result is a boost in blood O_2_ transport capacity that counteracts the reduced Hb-O_2_ affinity induced by the initial hypercapnia-induced acidosis.

Regulation of respiratory acidosis by sea bass exposed to hypercapnia in normal alkalinity sea water (TA ~2,800 μM) resulted in an elevation of plasma [HCO_3_−] by ~18 mM, which was correlated with a decrease in plasma [Cl-] of ~13 mM. In comparison, while we saw a slight rise in plasma [Na^+^] on initial exposure to hypercapnia, this was transient, and there was no overall correlation between plasma [Na^+^] and [H^+^] during the whole 135 minute experiment. However, a lack of increase in plasma [Na^+^] during acid-base regulation does not preclude increased Na^+^ uptake (to facilitate H_+_ excretion) during acid-base regulation. Instead, a lack of increased plasma [Na^+^] may simply reflect upregulation of the hypo-ionoregulatory mechanism for NaCl excretion in marine teleosts (Liu et al., 2016), which would presumably occur to compensate for enhanced uptake of Na^+^ to facilitate H^+^ excretion by NHE. This would also help explain the observed reduction in plasma [Cl^−^] in fish exposed to hypercapnia.

In comparison, sea bass exposed to hypercapnia in low alkalinity sea water (TA ~200 μM) showed no ability to accumulate HCO_3_−, to compensate for respiratory acidosis, and did not experience a decrease in plasma [Cl^−^]. At face value, these results may support potential direct uptake of HCO_3_− from seawater in exchange for blood Cl− through HCO_3_−/Cl− exchange across the gills (Esbaugh et al., 2012; Perry and Gilmour, 2006; Tovey and Brauner, 2018). However, the thermodynamics of this proposed mechanism are not clear, as [Cl^−^] is much higher in seawater than in internal fluids of fish, and the opposite is true for [HCO_3_−]. This implies that both counterions would have to be transported against their concentration gradients, and furthemore, these gradients would become increasingly unfavourable as blood acidosis is compensated. Importantly, our hypercapnic low alkalinity seawater had a pH of ~5.7, which was ~1.2 pH units lower than hypercapnic normal alkalinity seawater (a 15-fold increase in [H^+^]). Based on nominal values of [Na^+^] and [H^+^] inside fish gill ionocytes and calculations in Parks *et al.* (2008), the low alkalinity seawater would not sustain H^+^ excretion via NHEs (Table S5). Interestingly, as Na^+^ excretion is coupled to Cl^−^ excretion (via pathways independent of NHE), inactivation of NHE would also explain the lack of increase of plasma [Cl^−^] in low alkalinity sea water. Overall, this evidence supports enhanced NHE mediated H^+^ excretion (resulting in retention of metabolically produced HCO_3_− in the blood), rather than direct HCO_3_− uptake from sea water, as the primary mechanism underlying regulation of respiratory acidosis in sea bass.

To investigate the mechanisms used by sea bass to enhance acid-excretion, we examined whether changes in gill NKA and NHE3 occur after acute (~135 minute) exposure to hypercapnia. Gill NKA and NHE3 protein abundance did not change, ruling out increased protein synthesis as the mechanism responsible for the observed upregulation in acid-excretion; this is not surprising considering the short timeframe of our experiments. We also examined the potential translocation of pre-existing NKA and NHE3 to the ionocyte basolateral and apical membranes, respectively. Such mechanisms upregulate acid-base regulatory ion transport in elasmobranchs (Roa et al., 2014; Tresguerres et al., 2005; Tresguerres et al., 2006; Tresguerres et al., 2007b) and hagfish (Parks et al., 2007; Tresguerres et al., 2007a); however, NKA and NHE3 protein abundance in the gill membrane fraction of European sea bass gills was also unchanged, ruling out NKA and NHE3 translocation in our experiments. Finally, we hypothesized that sea bass could have remodelled the apical membrane of ionocytes to increase the sites for H^+^ excretion in contact with seawater. Indeed, this was the case as the surface area of the NHE3-abundant apical membrane of NKA-rich ionocytes roughly doubled in response to hypercapnia.

Previous studies on freshwater fishes have also documented morphological adjustments in gill ionocytes upon comparable hypercapnic exposures. However, the responses were the opposite to our study, as those freshwater fishes experienced a significant reduction in ionocyte apical surface area (Baker et al., 2009a; Goss et al., 1992b; Leino and McCormick, 1984). In some cases, the apical membrane retracted into a more pronounced apical pit (Goss et al., 1992a; Goss et al., 1994), which was suggested to create a microenvironment with higher [Na^+^] compared to the bulk freshwater and facilitate Na^+^/H^+^ exchange (Kumai and Perry, 2012). However, exposure to more pronounced hypercapnia (8 kPa CO_2_ over four days) induced an increase in gill ionocyte apical surface area in freshwater catfish (*Ictalurus punctatus*) (Cameron and Iwama, 1987). This response was similar to the seabass in our study; however, the longer time frame likely allowed for additional responses that were not investigated, such as increased synthesis of ion-transporting proteins or a change in the H^+^ excreting mechanism. In any case, the ability of seabass to rapidly compensate a blood respiratory acidosis by increasing gill ionocyte apical surface area is in large part possible due to the overabundance of Na^+^ in sea water, which establishes favourable conditions for NHE-mediated H^+^ excretion.

Freshwater species typically take from 24 h to > 72 h to regulate blood pH after exposure to 1 kPa CO_2_ (Baker et al., 2009a; Claiborne and Heisler, 1984; Claiborne and Heisler, 1986; Damsgaard et al., 2015; Larsen and Jensen, 1997; Perry, 1982; Perry et al., 1981; Smatresk and Cameron, 1982). While it is generally believed that marine teleosts can regulate their blood acid-base status at a faster rate than freshwater species (Brauner et al., 2019), relatively little research has been conducted to characterise the speed of acid-base regulation in marine fish. A bibliography search revealed four previous studies on five marine teleost species that characterized the time course of the acid-base regulatory response after exposure to 1 kPa CO_2_ (Fig. 8). Of these species, only the Japanese amberjack (*Seriola quinqueradiata*) was able to restore blood pH_e_ faster than sea bass (~60 min vs ~135 min; Fig. 8B). The remaining four species regulated blood pH_e_ between 3 and 24 h post CO_2_ exposure (Figure 8C, D, E, F).

**Figure 8:**
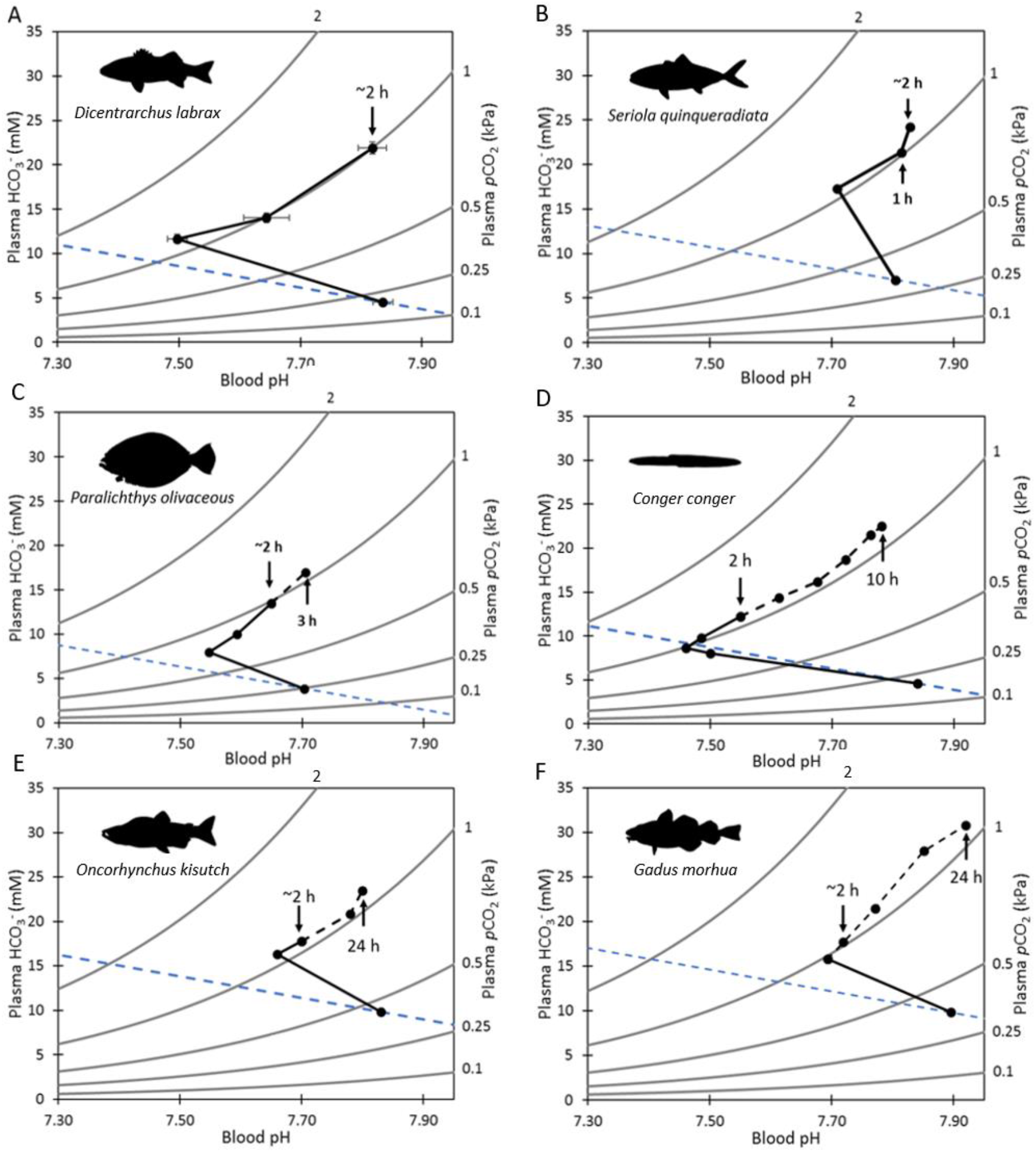
Blood pH/HCO_3_−/*p*CO_2_ plots for **A.** European sea bass, *Dicentrarchus labrax* (Present study), **B.** Japanese amberjack, *Seriola quinqeradiata* (Hayashi *et al.*, 2004, re-plotted raw data provided by pers. comm. with Dr Atsushi Ishimatsu, Can Tho University), **C.** Japanese flounder, *Paralichthys olivaceus* (Hayashi et al., 2004), **D.** conger eel, *Conger conger* (Toews et al., 1983), **E.** coho salmon, *Oncorhynchus kisutch* (Perry, 1982), and **F.** Atlantic cod, *Gadus morhua* (Larsen et al., 1997). The corresponding blood pH and HCO_3_− of each species at a time ~2 h after 1 kPa CO_2_ exposure is indicated to allow direct comparisons with European sea bass. Times below the relevant point indicate when blood pH was not statistically different from pre-exposure levels for each species. The time course of the acid-base response after 2 h is indicated by a dashed black line. The dashed blue line is an approximated non-HCO_3_− buffer line based on the mean haematocrit of sea bass from the present study and calculated using the equation for rainbow trout from Wood *et al.* (1982).

The robust ability of sea bass to rapidly acid-base regulate in response hypercapnia likely plays a significant role in their natural environment. Specifically, sea bass feed in shallow coastal estuaries and salt marsh habitats during the summer (Doyle et al., 2017), and these habitats typically experience large fluctuations in CO_2_ levels over short time periods (Hofmann et al., 2011; Melzner et al., 2013; Wallace et al., 2014). For example, equivalent salt marshes on the US coast regularly experience CO_2_ fluctuations of ~0.4 kPa across tide cycles during the summer (Baumann et al., 2015). The fast acid-base regulatory response observed in our study indicates that sea bass would be able to rapidly correct the respiratory blood acidosis caused by this level of CO_2_ variation in <1 hour. Critically, regulation of erythrocyte pH_i_ by sea bass was the fastest recorded in any fish species. By rapidly restoring erythrocyte pH_i_ O_2_ transport capacity is maintained, which is crucial for active predatory teleosts. However, environmental CO_2_ variation cannot be the sole driver for enhanced acid-base regulatory capacities in all species. For example, Japanese amberjack show a similarly fast blood acid-base regulatory response (Hayashi et al., 2004) but primarily inhabit pelagic, offshore ecosystems in which large variation in environmental CO_2_ may be less likely to occur. An alternative may be that active, predatory species have developed higher capacities for acid-base regulation to deal with large metabolic acidosis (as a result of anaerobic respiration used during intense exercise involved in prey capture). Understanding the mechanisms that determine species-specific differences in acid-base regulatory capacity will help understand differential impacts of acute exposure to elevated CO_2_, both by itself and in combination with other stressors such as hypoxia. For example, we have recently reported that sea bass showed enhanced hypoxia tolerance when exposed to progressive and environmentally relevant hypercapnia and hypoxia over a 6 hour period (Montgomery et al., 2019). In contrast, European plaice (*Pleuronectes platessa*) and European flounder (*Platichthys flesus*) exposed to the same conditions showed reduced hypoxia tolerance (P_crit_), which was associated with reduced Hb-O_2_ affinity and O_2_ uptake resulting from an uncompensated respiratory acidosis (Rogers, 2015; Montgomery *et al.* unpublished observations). Thus, species with more robust acid-base regulatory mechanisms seem more resilient to interactive effects between hypercapnia and hypoxia.

## Conclusion

Overall, our study highlights the capacity of European sea bass to rapidly (2 hours) regulate blood and erythrocyte acid-base status and O_2_ transport capacity upon exposure to a pronounced and sudden increase in environmental CO_2_ levels. Sea bass’ ability to rapidly upregulate H^+^ excretion appears to be mediated via the increased exposure of NHE3-containing apical surface area of gill ionocytes, rather than changes in NHE3 or NKA protein abundance or localisation. Additionally, sea bass erythrocyte pH_i_ is regulated even more rapidly than blood pH (40 minutes), which enables them to quickly restore the affinity of haemoglobin for O_2_, and therefore blood O_2_ transport capacity during exposure to elevated CO_2_. In conjunction, these acid-base regulatory responses will minimise the impact of pronounced and rapidly fluctuating CO_2_ in their natural environments, and so may prevent disruption of energetically costly activities such as foraging or digestion, and may make sea bass more resilient to impacts of hypoxia and additional stressors during acute periods of hypercapnia. This is an avenue where we believe further research effort is necessary.

## Conflict of Interest

The authors declare no conflicts of interest.

## Author contributions

D.W.M. designed the experiment, performed all data collection and analysis other than for gill samples, and wrote the manuscript. W.D., J.F., and A.B assisted with data collection, data analysis and editing of the manuscript. R.W.W. supervised the study and assisted with designing the experiments, data collection and analysis, and writing of the manuscript. S.D.S., G.H.E., and S.N.R.B. assisted with data analysis and editing of the manuscript. G.T.K. conducted all data analysis of gill samples and assisted with writing the manuscript. M.T. assisted with data analysis and writing of the manuscript.

## Acknowledgements

This work was supported by a NERC GW4+ Doctoral Training Partnership studentship from the Natural Environment Research Council [NE/L002434/1] with additional funding from CASE partner, The Centre for Environment, Fisheries and Aquaculture Science (Cefas) to D.W.M., and from the Biotechnology and Biological Sciences Research Council (BB/D005108/1 and BB/J00913X/1) and NERC (NE/H017402/1) to R.W.W. G.T.K. was funded by the National Science Foundation (NSF) Postdoctoral Fellowship Program (#1907334). M.T. received funding from NSF IOS #1754994. We thank Dr Junya Hiroi for providing the NHE3 antibodies, and Dr Andrew Esbaugh and Dr Till Harter for discussions related to data interpretation. We would similarly like to thank the aquarium staff, particularly Dr Gregory Paul, Paul Tyson, Rebecca Turner, Alex Bell, and Richard Silcox, of the Aquatic Resource Centre at the University of Exeter for assistance with fish husbandry and maintenance of aquarium facilities.

## Data availability

Data will be deposited on the University of Exeter’s Open Research Exeter (ORE) depository and a URL to the dataset provided prior to article acceptance or upon request by reviewers.

## 1. Supplementary materials

**Figure S1:**
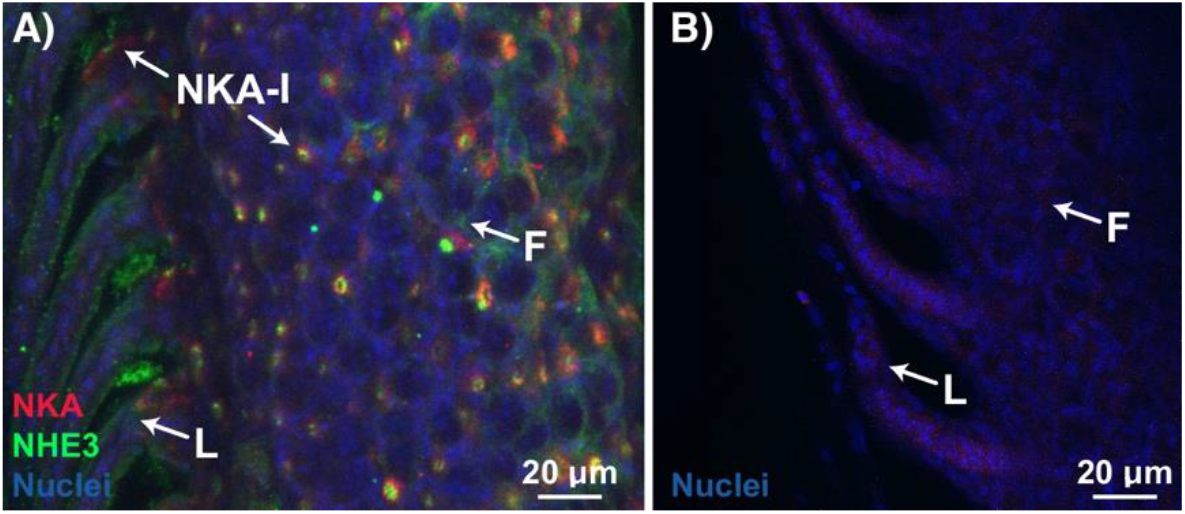
**A.** Na^+^/K^+^-ATPase (NKA, red) and Na^+^/H^+^ Exchanger 3 (NHE3, green) immunostaining within European sea bass gill. Ionocytes containing NKA and NHE3 (NKA-I) were observed on the filament (F) and base of lamellae (L). **B.** Negative controls (no primary antibodies) had no discernible signal. Nuclei (blue) are stained with DAPI.

**Table S1:** Water chemistry of gill irrigation chambers used while blood sampling fish

|  | Exposure length |  |  |  |
| --- | --- | --- | --- | --- |
|  | 0 min | ~10 min | ~40 min | ~135 min |
| Temperature ($^{\circ}\text{C}$) | $14.00 \pm 0.00$ | $14.05 \pm 0.15$ | $14.00 \pm 0.30$ | $13.95 \pm 0.15$ |
| pH (NBS) | $8.14 \pm 0.02$ | $6.95 \pm 0.02$ | $6.91 \pm 0.01$ | $6.91 \pm 0.04$ |
| Salinity | $34.6 \pm 0.4$ | $35.6 \pm 0.5$ | $34.8 \pm 0.7$ | $34.6 \pm 0.7$ |
| $p\text{CO}_2$ ($\mu\text{atm}$) | $0.060 \pm 0.010$ | $0.965 \pm 0.045$ | $1.227 \pm 0.164$ | $1.024 \pm 0.088$ |
| TA ($\mu\text{M}$) | $3120 \pm 372$ | $2834 \pm 36$ | $3301 \pm 361$ | $2750 \pm 27$ |

**Table S2:**
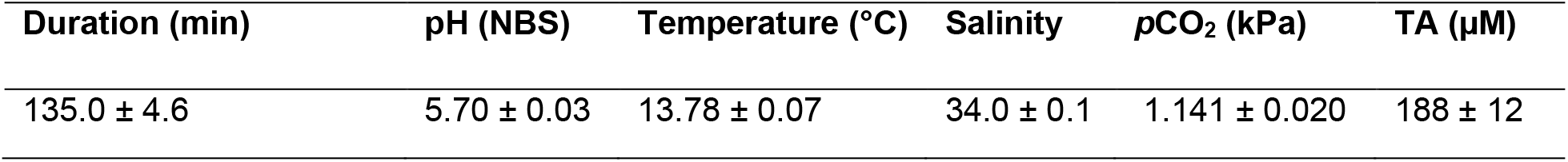
Mean ± s.e.m. of water chemistry parameters within isolation tanks during low alkalinity hypercapnia exposure

| Duration (min) | pH (NBS) | Temperature ($^{\circ}\text{C}$) | Salinity | $p\text{CO}_2$ (kPa) | TA ($\mu\text{M}$) |
| --- | --- | --- | --- | --- | --- |
| $135.0 \pm 4.6$ | $5.70 \pm 0.03$ | $13.78 \pm 0.07$ | $34.0 \pm 0.1$ | $1.141 \pm 0.020$ | $188 \pm 12$ |

**Table S3:** Water chemistry of gill irrigation chambers used while blood sampling fish in low alkalinity water

| pH (NBS) | Temperature ($^{\circ}\text{C}$) | Salinity | $p\text{CO}_2$ ($\mu\text{atm}$) | TA ($\mu\text{M}$) |
| --- | --- | --- | --- | --- |
| $5.53 \pm 0.31$ | $14.05 \pm 0.05$ | $35.05 \pm 0.25$ | $1.288 \pm 0.104$ | $188 \pm 129$ |

**Table S4:** Mean ± s.e.m. of water chemistry parameters within isolation boxes prior to gill sampling. Gill samples from sea bass exposed to hypercapnia were taken from 5 sea bass immediately after flux measurements were completed.

| Treatment | Duration (min) | pH (NBS) | Temperature (°C) | Salinity | $p\text{CO}_2$ (kPa) | TA ($\mu\text{M}$) |
| --- | --- | --- | --- | --- | --- | --- |
| Normocapnia | n/a | $8.05 \pm 0.00$ | $14.12 \pm 0.06$ | $33.6 \pm 0.0$ | $0.055 \pm 0.001$ | $2335 \pm 7$ |
| Hypercapnia | $132.9 \pm 2.6$ | $6.97 \pm 0.02$ | $14.14 \pm 0.02$ | $33.5 \pm 0.0$ | $0.770 \pm 0.037$ | $2365 \pm 9$ |

**Table S5:**
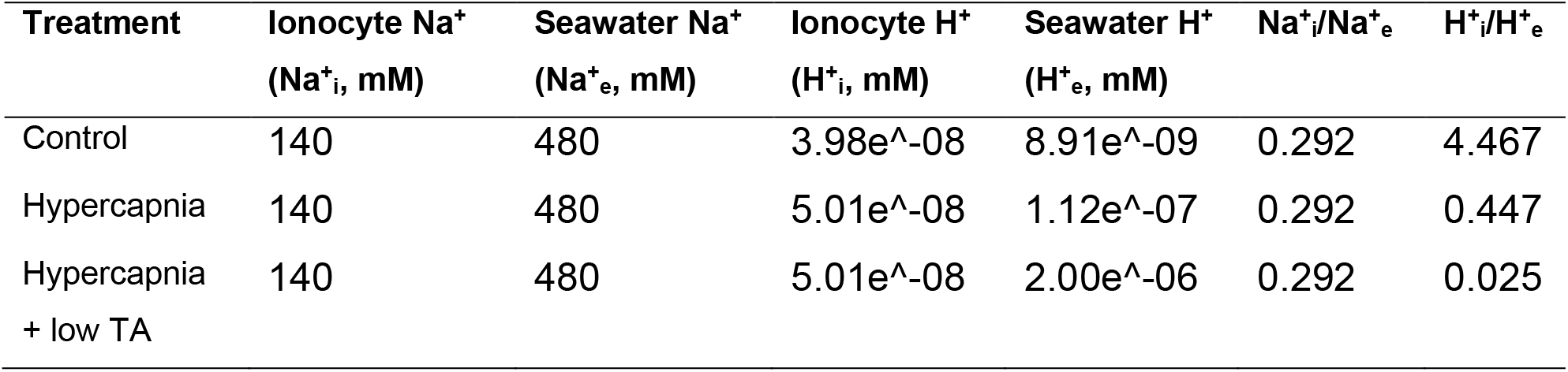
Theoretical calculations of H^+^ excretion by NHE in response to environmental hypercapnia. Calculations based on Parks *et al.* 2008. If Na^+^_i_/Na^+^_e_ > H^+^_i_/H^+^_e_ then H^+^ excretion by NHE is thermodynamically unviable.

| Treatment | Ionocyte $\text{Na}^+$<br>($\text{Na}^+_{\text{i}}$, mM) | Seawater $\text{Na}^+$<br>($\text{Na}^+_{\text{e}}$, mM) | Ionocyte $\text{H}^+$<br>($\text{H}^+_{\text{i}}$, mM) | Seawater $\text{H}^+$<br>($\text{H}^+_{\text{e}}$, mM) | $\text{Na}^+_{\text{i}}/\text{Na}^+_{\text{e}}$ | $\text{H}^+_{\text{i}}/\text{H}^+_{\text{e}}$ |
| --- | --- | --- | --- | --- | --- | --- |
| Control | 140 | 480 | $3.98\text{e}^{-08}$ | $8.91\text{e}^{-09}$ | 0.292 | 4.467 |
| Hypercapnia | 140 | 480 | $5.01\text{e}^{-08}$ | $1.12\text{e}^{-07}$ | 0.292 | 0.447 |
| Hypercapnia<br>+ low TA | 140 | 480 | $5.01\text{e}^{-08}$ | $2.00\text{e}^{-06}$ | 0.292 | 0.025 |

## Notes

### Competing Interest Statement

The authors have declared no competing interest.

